# Characterization of the gastric motility response to human motilin and erythromycin in human motilin receptor-expressing transgenic mice

**DOI:** 10.1101/436436

**Authors:** Shinichi Kato, Aoi Takahashi, Mai Shindo, Ayano Yoshida, Tomoe Kawamura, Kenjiro Matsumoto, Bunzo Matsuura

## Abstract

Motilin is a gastrointestinal peptide hormone that stimulates gastrointestinal motility. Motilin is produced primarily in the duodenum and jejunum. Motilin receptors (MTLRs) are G protein-coupled receptors that may represent a clinically useful pharmacological target as they can be activated by erythromycin. The functions of motilin are highly species-dependent and remain poorly understood. As a functional motilin system is absent in rodents such as rats and mice, these species are not commonly used for basic studies. In this study, we examine the usefulness of human MTLR-overexpressing transgenic (hMTLR-Tg) mice by identifying the mechanisms of the gastric motor response to human motilin and erythromycin.

The distribution of hMTLR was examined immunohistochemically in male wild-type (WT) and hMTLR-Tg mice. The contractile response of gastric strips was measured isometrically in an organ bath, while gastric emptying was determined using phenol red.

hMTLR expression was abundant in the gastric smooth muscle layer but more potently expressed in the myenteric plexus of hMTLR-Tg mice but not WT mice. hMTLR was not co-localized with vesicular acetylcholine transporter, a marker of cholinergic neurons in the myenteric plexus. Treatment with human motilin and erythromycin caused concentration-dependent contraction of gastric strips obtained from hMTLR-Tg mice but not from WT mice.

The contractile response to human motilin and erythromycin in hMTLR-Tg mice was affected by neither atropine nor tetrodotoxin and was totally absent in Ca^2+^-free conditions. Furthermore, intraperitoneal injection of erythromycin significantly promoted gastric emptying in hMTLR-Tg mice but not in WT mice.

Human motilin and erythromycin stimulate the gastric motor response in hMTLR-Tg mice. This action is mediated by direct contraction of smooth muscle via the influx of extracellular Ca^2+^. Thus, hMTLR-Tg mice may be useful for the evaluation of MTLR agonists as gastric prokinetic agents.

## Introduction

Motilin, a 22-amino acid peptide, was firstly identified in 1973 from porcine intestine [1]. The most important physiological function of this hormone is the stimulation of gastrointestinal motility. Motilin is thought to be primarily involved in gastric phase III activity of the migrating motor contraction (MMC) observed during the fasting state. Indeed, several studies have shown that plasma motilin concentrations are increased in parallel with the contractile response in the stomach [2, 3]. In addition, exogenous administration of motilin caused MMC phase III-like contractions, while gastric phase III contractions were inhibited by anti-motilin serum and administration of a motilin antagonist [4–6].

The receptor for motilin (MTLR) was identified in 1999 from human stomachs and was previously known as GPR38 [7]. MTLR is a G protein-coupled receptor and has 52% sequence homology with ghrelin receptors [8]. Previous studies have demonstrated that MTLR is expressed in gastrointestinal muscle cells and the myenteric plexus, but not in the mucosa and submucosa of the gastrointestinal tract in humans [9, 10]. These findings are consistent with results that suggest that the motilin-induced motor response is mediated through direct action on smooth muscle cells and the activation of cholinergic pathways [11–13]. Furthermore, the antibiotic drug erythromycin has been shown to stimulate MTLRs and mimic the motor response observed following treatment of motilin [14, 15].

The actions of motilin are known to be highly species-dependent. In addition to humans, pigs, and dogs, motilin has been identified in rabbits, cows, cats, sheep, horses and shrews; in contrast, a functional motilin system is absent in rodents such as mice, rats, and guinea pigs [16–22]. A pseudogene of MTLR has been identified in rodents that fails to respond to motilin, which may be a result of differences in the anatomy and physiology of rodents; this is corroborated by the inability of these organisms to vomit [20, 21]. Further, since these rodents are commonly used for basic science studies of many pathways, the lack of a motilin system prevents their use for elucidating the basic physiological and pathophysiological functions of motilin. It also means these models are not useful for the screening of MTLR agonists.

We have previously produced human MTLR-overexpressing transgenic (hMTLR-Tg) mice. hMTLR-Tg mice had an increased intake of food and water during human motilin and erythromycin treatment when compared to wild-type (WT) mice [23]. However, the effect of human motilin and erythromycin treatment on gastric motor activity in hMTLR-Tg mice is yet to be examined.

In order to identify the usefulness of hMTLR-Tg mice, in this study, we examined and characterized the gastric motor response following human motilin and erythromycin treatment in hMTLR-Tg mice.

## Materials and methods

### Animals and ethics statement

This study was carried out in strict accordance with the recommendations found in the Guide for Care and Use of Laboratory Animals (8^th^ Edition, National Institutes of Health). The protocols were approved by the committee on the Ethics of Animal Research of the Kyoto Pharmaceutical University (Permit Number: 16-014). Male C57BL/6 mice (8–10 weeks) weighing 22–26 g were purchased from Japan SLC Inc. (Shizuoka, Japan). Human motilin receptor-overexpressing transgenic (hMTLR-Tg) mice were generated on an C57BL/6 mice background as described previously [23]. All mice were maintained in plastic cages with free access to food and water. All animals were housed at 22 ± 1 °C with a 12-hour light/dark cycle.

### Drugs

Human motilin was purchased from Anygen Co., Ltd (GwangJu, Korea). Erythromycin was purchased from Mylan (Tokyo, Japan). Acetylcholine (ACh), atropine, and tetrodotoxin (TTX) were purchased from Wako Pure Chemical (Osaka, Japan). For *in vivo* experiments (gastric emptying), erythromycin was diluted in physiological salt solution. The dose of erythromycin was expressed as mg/kg body weight. For *in vitro* experiments (contraction), erythromycin was diluted in, and other drugs were dissolved in, physiological salt solution. Drug concentrations were expressed as final molar concentrations in the organ bath solution. All drugs were prepared immediately before use.

### Immunohistochemistry

Tissue preparation and immunohistochemical procedures were performed as described previously [24, 25]. After 18 hours of fasting, the stomach was removed. For glass-mounted sections, tissues were fixed by immersion in 4% paraformaldehyde (PFA; in 0.1 M phosphate buffer) for 2 hours at 4°C, cryoprotected in 0.1 M 20% sucrose (in phosphate buffer) overnight, and subsequently frozen in OCT compound (Sakura Finetec, Tokyo, Japan). The preparation was then cut into 30 μm sections using a cryostat (CM3050, Leica Biosystems, Nussloch, Germany) and mounted onto Superfrost Plus slides (Matsunami Glass, Osaka, Japan). For whole-mounted sections, the mucosal and submucosal layers were gently removed under a dissecting microscope and the muscularis preparations were fixed in 4% PFA (in 0.1 M phosphate buffer) for 30 min at 4°C, and cryoprotected in 0.1 M 20% sucrose in phosphate buffer overnight. Glass-mounted and whole mounted sections were treated with 10% donkey serum containing 0.2% Triton X-100, in phosphate buffered saline (PBS), for 1 hour. Subsequently, the preparations were probed for 40 hours at room temperature with rabbit anti-human motilin antibody (MBL, Aichi, Japan), goat anti-vesicular acetylcholine transporter (VAChT) (Phonix Pharmaceuticals, Burlingame, CA, USA), and Alexa Flour 488-conjugated mouse anti-alpha smooth muscle actin (α-SMA) antibody (Abcam, Cambridge, MA, USA). After washing in PBS, preparations were labeled for 4 h at room temperature with donkey anti-rabbit IgG (conjugated to Alexa Fluor 488 or 594) and anti-goat IgG (conjugated to Alexa Fluor 594) (1:400, Thermo Fisher Scientific, Waltham, MA, USA). Preparations were imaged on a confocal microscope (A1R+, Nikon, Tokyo, Japan) using NIS-Elements AR 4.20,00 software (Nikon) to capture images, project and reconstruct multiple images in Z-stacks.

### Determination of gastric contractile response in vitro

After 18 hours of fasting, the stomach was removed and immediately placed in a freshly prepared normal physiological salt solution (136.9 mM NaCl, 5 mM KCl, 1.5 mM CaCl_2_, 1 mM MgCl_2_, 23.8 mM NaHCO_3_, 5.5 mM glucose, and 0.01 mM ethylene-diamine tetraacetic acid (EDTA); pH 7.4). Ca^2+^-free solutions were prepared by removing CaCl_2_ and adding 0.5 mM EGTA in a normal physiological salt solution. Muscle strips (3×5 mm) were obtained from the fundus, corpus and antrum regions along the circular axis, and then suspended in an organ bath filled with physiological salt solution at 37°C in an atmosphere of 95% O_2_:5% CO_2_. Contractile responses were measured isometrically under a resting tension of 10 mN and this data recorded (Pantos Co., Ltd, Kyoto, Japan).

### Determination of gastric emptying in vivo

Gastric emptying was determined using the phenol red method described previously [26]. Briefly, after 18 hours of fasting, mice received 1.5% carboxymethlcellulose (CMC, Nacali Tesque, Kyoto, Japan) containing 0.05% phenol red (Wako Pure Chemical) orally at a volume of 0.5 mL/mouse. Ten minutes later, mice were sacrificed by cervical dislocation, their abdomen were opened, and the gastroesophageal junction and pylorus were clamped. Then, the stomach was removed, placed in 0.1 M NaOH and homogenized. The suspension was allowed to settle for 1 hour at room temperature, after which 2% trichloroacetic acid (final concentration) was added and samples were centrifuged at 2,500 g for 10 min. The supernatant was mixed with 0.05 M NaOH, and the amount of phenol red was determined colorimetrically at 560 nm using a microplate reader (Multiskan GO, Thermo Fisher). The standard sample (0%) was determined by the amount of phenol red recovered from mice sacrificed immediately after oral administration of 1.5% CMC containing 0.05% phenol red. Gastric emptying rate was calculated according to the following equation;

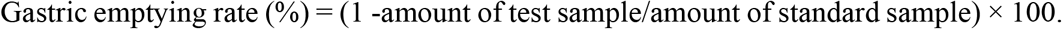

### Statistical analysis

Data are presented as mean ± SEM. Statistical analysis was performed with GraphPad Prism 6.0h (GraphPad Software, La Jolla, CA, USA) using a student t-test and one-way or two-way ANOVA followed by Bonferroni’s test. We considered p values less than 0.05 to represent statistically significant results.

## Results

### Distribution of hMTLR in the stomach

To confirm the presence of hMTLR in the stomach of hMTLR-Tg mice, immunohistochemical analysis was performed hMTLR-Tg and WT mice using an anti-hMTLR antibody. In glass-mounted sections, hMTLR-positive immunostaining was not detected in the stomach obtained from WT mice (Fig 1A and B). However, in contrast, hMTLR-positive immunostaining was observed both in the muscle layer and more potently in the myenteric plexus of stomachs obtained from hMTLR-Tg mice. Potent expression of hMTLR was also found in gastric epithelial cells of hMTLR-Tg mice. To characterize the localization of hMTLR, double immunostaining using an anti-hMTLR antibody with an anti-VAChT antibody (a marker of cholinergic neurons) and an anti-α-SMA (a marker of smooth muscle) was performed in the stomach of WT and hMTLR-Tg mice. VAChT staining was detected in neuronal fibers of the smooth muscle layer and myenteric plexus of both WT and hMTLR-Tg mice (Fig 1A). The expression of VAChT in the myenteric plexus was mostly colocalized with hMTLR-positive staining except in neuronal fibers. Positive immunostaining for α-SMA was also identified in the smooth muscle layer of both WT and hMTLR-Tg mice (Fig 1B). Double immunostaining revealed the colocalization of α-SMA and hMTLR in muscle layer of hMTLR-Tg mice.

To further investigate the expression of hMTLR in the myenteric plexuses of hMTLR-Tg mice, double immunostaining of anti-hMTLR antibody was performed. In these experiments, alongside an anti-MTLR antibody, anti-VAChT antibody and anti-α-SMA antibodies were used in whole mounted sections of the stomach from hMTLR-Tg mice. Although the colocalization of hMTLR and α-SMA was observed in the myenteric plexus, hMTLR-positive staining was not consistent with VAChT-positive staining in the myenteric plexus.

**Fig 1.**
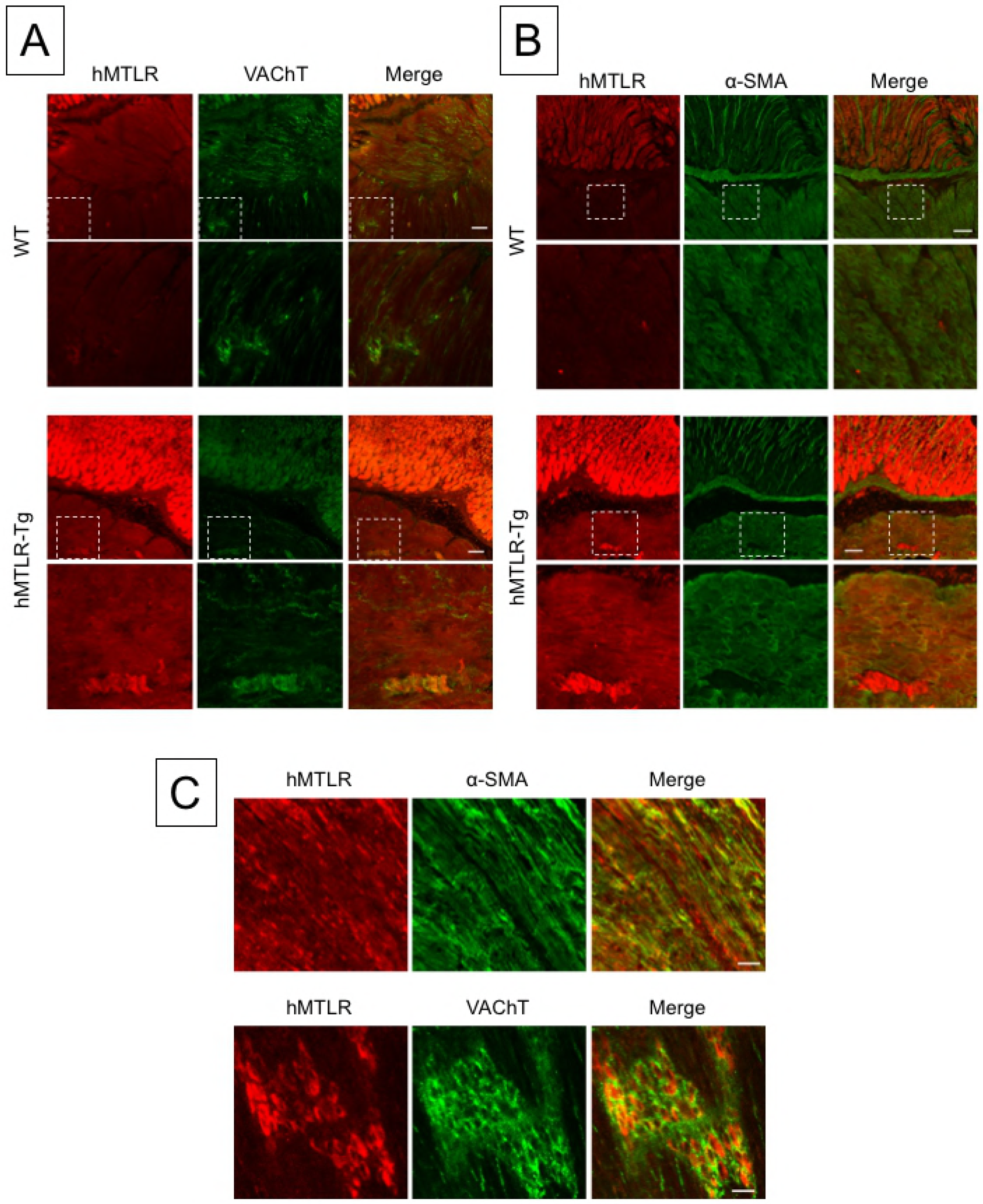
Distribution of hMTLR expression in the stomach of WT and hMTLR-Tg mice. (A) Double staining of hMTLR (red) with VAChT (green), a marker of cholinergic neurons, in the stomach of WT and hMTLR-Tg mice. Scale bar = 50 μm. (B) Double staining of hMTLR (red) with α-SMA (green), a marker of smooth muscles, in the stomach of WT and hMTLR-Tg mice. Scale bar = 50 μm. (C) Double staining of hMTLR (red) with α-SMA (green) or VAChT (green) in the whole mounted section (×600). Scale bar = 20 μm. The lower panel in each group indicates higher magnification of the square in the upper panels.

### Gastric contractile response to human motilin and erythromycin

To examine the gastric contractile response to human motilin, human motilin (10^−10^–10^−6^ M) and erythromycin (10^−9^–10^−5^ M) were introduced into an organ bath and the contractile amplitude of the gastric fundus, corpus and antrum obtained from WT and hMTLR-Tg mice. Although human motilin had no influence on smooth muscle tension in WT mice, motilin caused contractile responses at all sites examined in a concentration-dependent manner (Fig 2A and B). The order of the contractile response to human motilin was fundus > corpus > antrum. Likewise, erythromycin caused a concentration-dependent contractile response at all sites obtained from hMTLR-Tg mice but not WT mice (Fig 3A and B). The order of the contractile response to erythromycin was also fundus > corpus > antrum. The contractile response of gastric strips was more potent to human motilin than erythromycin.

**Fig 2.**
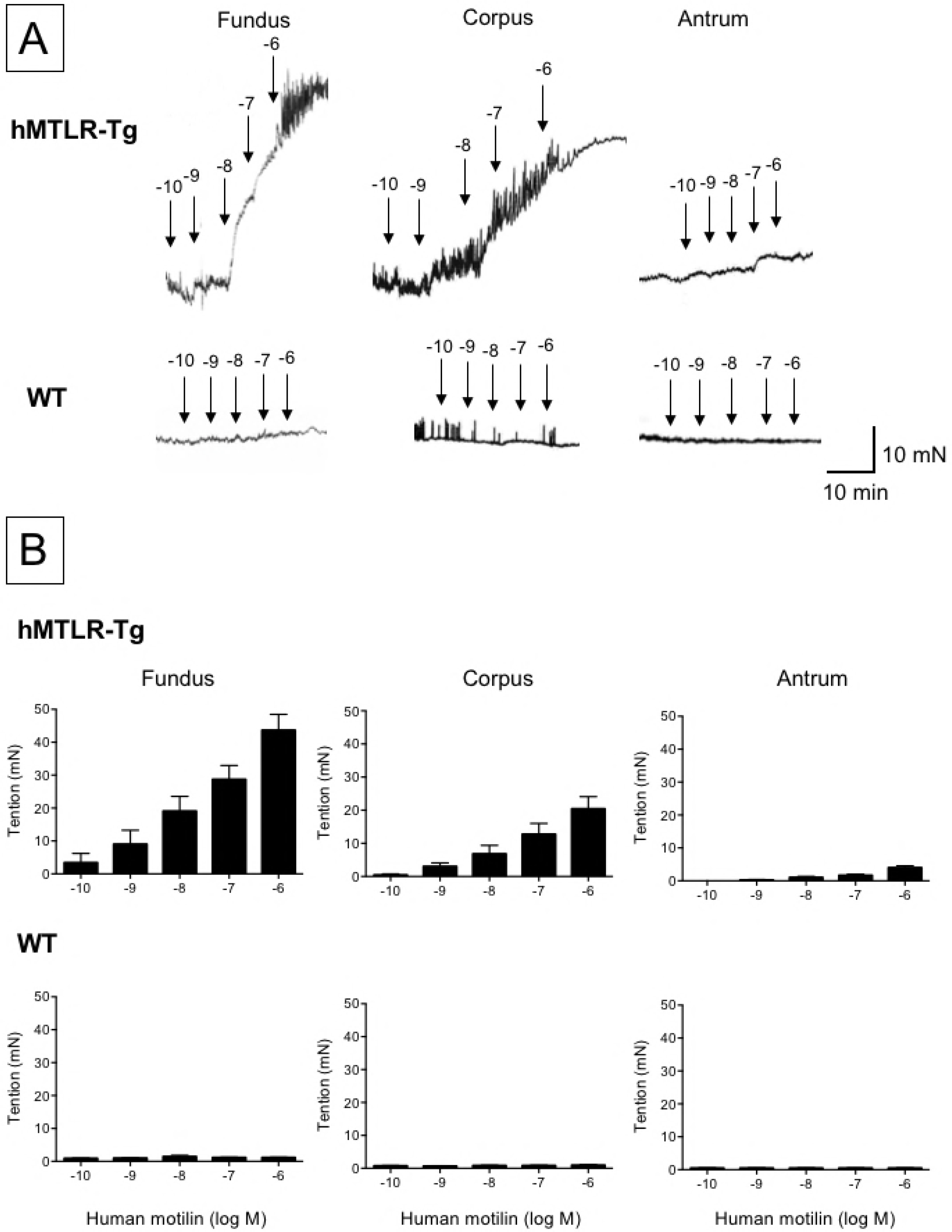
Effect of human motilin on smooth muscle tension of the gastric fundus, corpus and antrum from WT and hMTLR-Tg mice. (A) Typical recording and (B) summary of contractile response to human motilin (10^−10^–10^−6^ M) in the gastric fundus, corpus and antrum obtained from hMTLR-Tg (upper) and WT mice (lower). Data are presented as the mean ± SEM for 5 animals.

**Fig 3.**
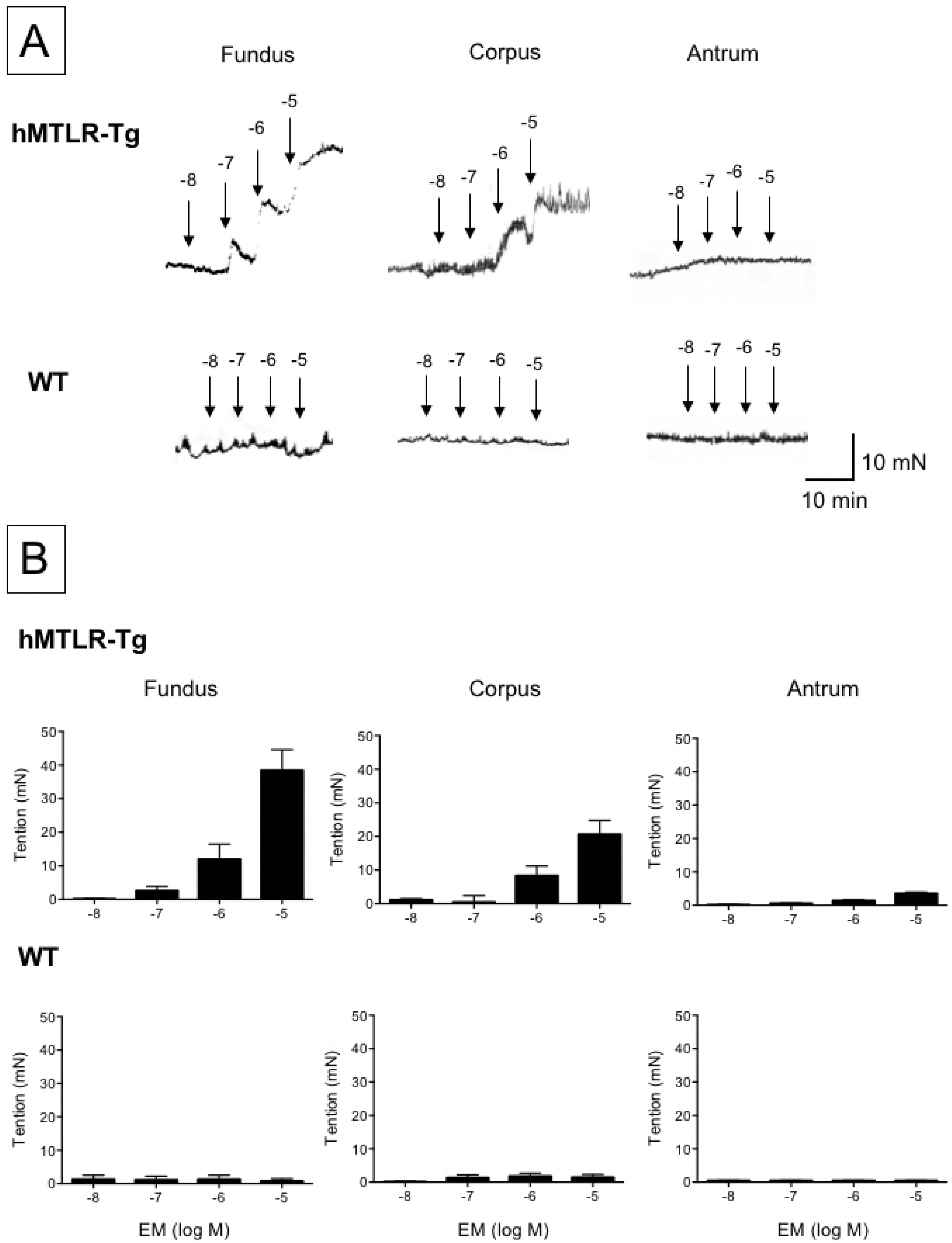
Effect of erythromycin on smooth muscle tension of gastric fundus, corpus and antrum obtained from WT and hMTLR-Tg mice. (A) Typical recording and (B) summary of the contractile response to human motilin (10^−9^–10^−5^ M) in gastric fundus, corpus and antrum obtained from hMTLR-Tg (upper) and WT mice (lower). Data are presented as the mean ± SEM for 5 animals.

### Gastric contractile response to acetylcholine

To confirm the specificity of the gastric contractile response to human motilin and erythromycin in hMTLR-Tg mice, the gastric contractile response to acetylcholine was examined in WT and hMTLR-Tg mice. Acetylcholine (10^−5^ M) caused visible contractile responses in the gastric fundus, corpus and antrum obtained from both WT and hMTLR-Tg mice (Fig 4). The gastric contractile responses in WT and hMTLR-Tg mice were almost similar. In this case, the order for contractile responses were: fundus > corpus > antrum. The contractile response was totally abolished by co-treatment with atropine (10^−5^ M).

**Fig 4.**
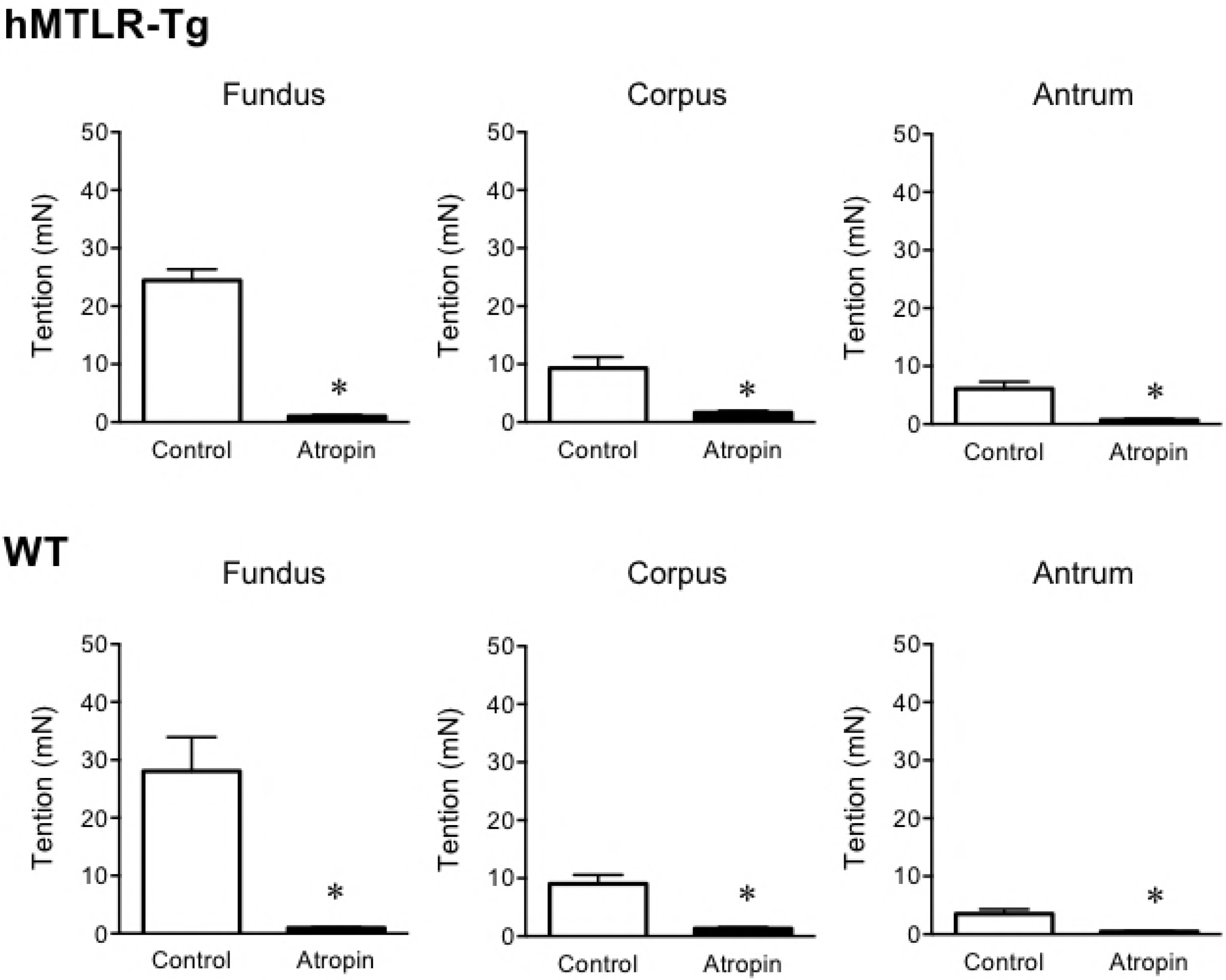
Effect of acetylcholine on smooth muscle tension of the gastric fundus, corpus and antrum obtained from WT and hMTLR-Tg mice. A summary of contractile responses to acetylcholine (10^−9^–10^−5^ M) with or without atropine (10^−6^ M) in the gastric fundus, corpus and antrum obtained from hMTLR-Tg (upper) and WT mice (lower) is shown. Data are presented as mean ± SEM for 5 animals. *Statistically significant difference from control (vehicle alone) at P < 0.05.

### Characterization of gastric contractile response to human motilin and erythromycin in hMTLR-Tg mice

We next sought to characterize the gastric contractile response to human motilin and erythromycin in hMTLR-Tg mice. The effect of atropine, tetrodotoxin and Ca^2+^-free conditions on motilin and erythromycin-dependent contractile responses in the gastric corpus obtained from hMTLR-Tg mice were examined. Human motilin (10^−6^ M) caused visible contractile response in the gastric fundus of hMTLR-Tg mice (Fig 5A and B). This contractile response was diminished by co-treatment with atropine (10^−6^ M) or tetrodotoxin (10^−6^ M). However, the contractile response was completed abrogated in Ca^2+^-free conditions. Likewise, the erythromycin-induced contractile response was not affected by atropine or tetrodotoxin, but totally abolished in Ca^2+^-free conditions.

**Fig 5.**
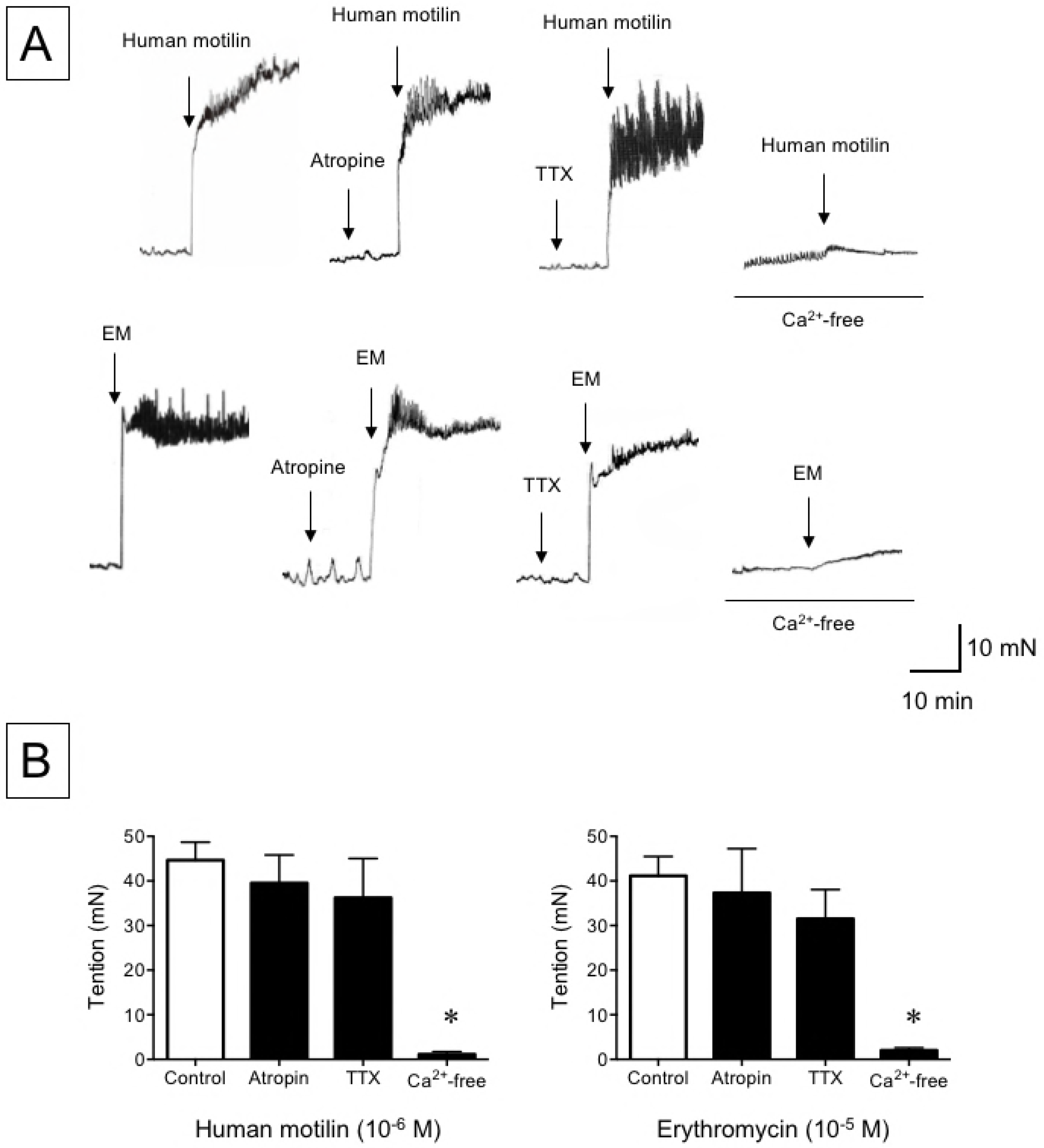
Effect of atropine, tetrodotoxin and Ca^2+^-free conditions on the contractile response to human motilin and erythromycin in the gastric fundus of hMTLR-Tg mice. (A) Typical recording and (B) summary of the contractile response to human motilin (10^−6^ M) (upper) and erythromycin (EM; 10^−5^ M) (lower) in gastric fundus obtained from hMTLR-Tg mice. Atropine (10^−6^ M) and tetrodotoxin (TTX; 10^-6^ M) were pretreated before the application of either human motilin or erythromycin (EM). A Ca^2+^-free solution was prepared by removing CaCl_2_ and adding 0.5 mM EGTA in an otherwise normal physiological salt solution. Data are presented as mean ± SEM for 5 animals. * Statistically significant difference from control (vehicle alone) at P < 0.05.

### Effect of erythromycin on gastric emptying in WT and hMTLR-Tg mice

Finally, we examined the effect of erythromycin on gastric emptying in hMTLR-Tg mice *in vivo*. Gastric emptying rates in WT and hMTLR-Tg mice were 53.5 ± 7.8% and 54.4 ± 5.1% at 10 min after oral administration of 1.5% CMC containing 0.05% phenol red, respectively (Fig 6). There were no significant differences in gastric emptying between WT and hMTLR-Tg mice. Intraperitoneal injection of erythromycin (10 mg/kg) had no effect on gastric emptying rate in WT mice (53.2 ± 5.9%) but significantly promoted gastric emptying rate to 70.3 ± 4.4% in hMTLR-Tg mice.

**Fig 6.**
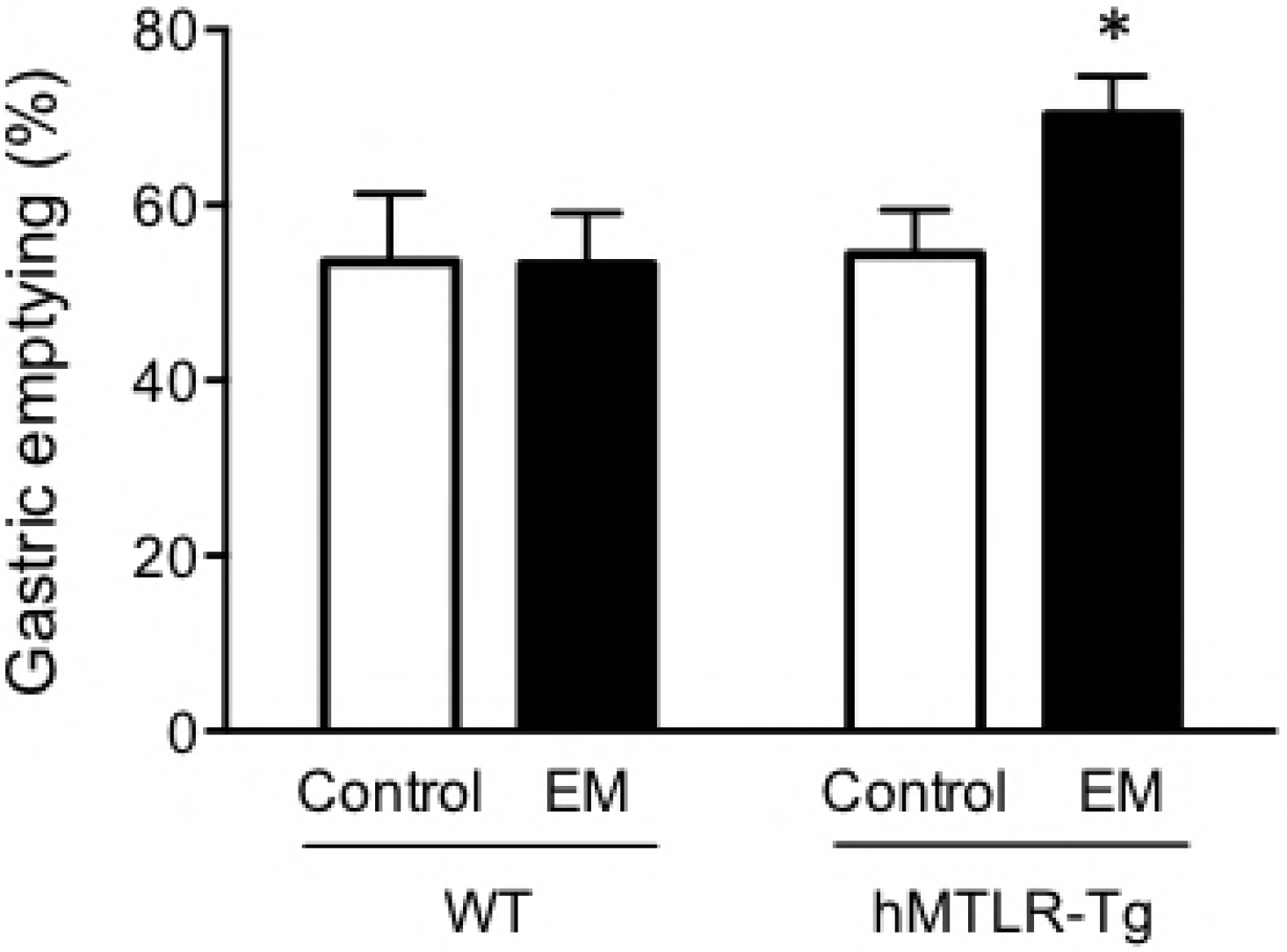
Effect of erythromycin on gastric emptying in WT and hMTLR-Tg mice. Animals were orally administered phenol red-containing CMC and the concentration of phenol red in the gastric content was determined 10 min later. Erythromycin (EM; 10 mg/kg) was injected intraperitoneally 20 minutes before phenol red administration. Data are presented as the mean ± SEM for 6 animals. *Statistically significant difference from control (vehicle alone) at P < 0.05.

## Discussion

Motilin is a peptide hormone that stimulates gastrointestinal motor activity via actions at MTLR receptors. Rodents, such as rats and mice, fail to respond to motilin due to lack of functional a signaling pathway and MTLR receptors [20, 21]. This fact impedes progress of motilin research because these rodents are commonly used for other basic scientific studies. We have previously produced an hMTLR-Tg mouse that has hMTLR cDNA incorporated into the transgene; this cDNA is designed to direct expression ubiquitously in mice [23]. In this study, we examined the expression of hMTLR and the observed motor activity in response to human motilin and erythromycin in hMTLR-Tg mice.

Immunohistochemical analyses identified enhanced expression of hMTLR in the smooth muscle layer, myenteric plexus and stomach mucosa of hMTLR-Tg mice, but not WT mice. Takeshita *et al.* [10] reported that mRNA expression of MTLR were detectable in the human gastrointestinal tract. Further, they showed MTLR-immunoreactivity in the smooth muscle layer and myenteric plexus, but not the mucosa and submucosa. A similar result was reported by Ter Beek *et al.* [27], although in this instance, they found expression of MTLR even in the mucosa. Although the expression of MTLR in the mucosa in basic studies is controversial, the expression of MTLR in the smooth muscle layer and myenteric plexus of hMTLR-Tg mice are similar to that in humans. We further observed that hMTLR in the myenteric plexus was not colocalized with VAChT, a maker of cholinergic neurons. Several studies have demonstrated that the gastrointestinal motor response to motilin is caused by the direct action of this agent on smooth muscle cells and the activation of cholinergic pathways as evidenced in rabbits and shrew, as well as humans [11–13, 28, 29]. These findings may suggest that a lack of motilin-induced motor response in hMTLR-Tg mice via activation of the cholinergic pathway occurs in contrast to the mechanism observed in humans.

The most important finding in this study was that human motilin produced concentration-dependent contractile responses in gastric smooth muscles strips obtained from hMTLR-Tg mice. Of course, human motilin failed to induce contraction in gastric smooth muscles obtained from WT mice. In addition, we show that the order of the contractile response to motilin was gastric fundus < corpus < antrum. An especially apparent contractile response was observed even at a low concentration (10^−10^ M) of human motilin in the fundus, but not corpus and antrum. Similar results have been reported in shrew and human stomachs [29, 30], suggesting that the motilin-induced contractile response is anatomically region-specific.

Similar to human motilin, erythromycin produced a concentration-dependent gastric contractile response, and this response was more pronounced in fundi obtained from hMTLR-Tg mice. However, the contractile response to erythromycin was about one tenth of that to human motilin, as consistent with a previous report in the human stomach [31]. These findings strongly suggest that the gastric contractile response to human motilin and erythromycin in hMTLR-Tg mice closely resembles the response observed in humans.

In the present study, we observed that acetylcholine produced contractile responses not only in hMTLR-Tg mice, but also in WT mice. Further, the order of contractile response to acetylcholine was similar to that seen with human motilin and erythromycin: gastric fundus > corpus > antrum. Thus, there is no difference in the contractile ability of smooth muscles between hMTLR-Tg and WT mice.

Interestingly, the acetylcholine muscarinic receptor antagonist atropine, and the neurotoxin tetrodotoxin, failed to attenuate the contractile responses to human motilin and erythromycin in hMTLR-Tg mice. Several studies have demonstrated that motilin-induced motor responses are mediated through direct action on smooth muscles and the activation of cholinergic pathways in dogs, rabbits, shrew and humans [11–13, 32]. In contrast, Satoh *et al.* [31] showed that the contractile response induced by an erythromycin-derivative in human gastric antrum was not influenced by atropine and tetrodotoxin. Further, in human and rabbit isolated stomachs, cholinergic contraction was caused by low concentrations of motilin and erythromycin, while the higher concentrations directly contracted the muscle [12, 33–35]. Thus, it is likely that the gastric contractile response to motilin and erythromycin may be mediated by the different pathways (i.e. either cholinergically or directly stimulated) depending on the concentration of each agent. As mentioned above, while the present study revealed that hMTLR was expressed in the myenteric plexus in addition to smooth muscles, the expression of hMTLR in the myenteric plexus was not colocalized with cholinergic neurons in hMTLR-Tg mice. Thus, the observations of contractile response to human motilin and erythromycin independent of cholinergic pathways fit well with the results obtained from immunohistochemical analyses. These findings suggest that hMTLR-Tg mice reproduce motilin-induced motor activity by direct action on smooth muscle cells and not via the operation of cholinergic pathways.

Several studies have demonstrated that the contractile effect of motilin and erythromycin is mediated via intracellular Ca^2+^ signaling in smooth muscle of rabbits, dogs and humans [36–39]. Further, Van Assche *et al.* [39] showed that motilin-induced increases in intracellular Ca^2+^ levels are dependent on Ca^2+^ influx; this is in contrast to acetylcholine, which triggers the release from intracellular Ca^2+^ stores. In this study, we observed that the gastric contractile response to human motilin and erythromycin in hMTLR-Tg mice was totally abolished in Ca^2+^-free conditions. Thus, it is likely that human motilin and erythromycin can produce direct contraction of gastric smooth muscle cells in hMTLR-Tg mice through activation of Ca^2+^ influx via MTLR.

Finally, we examined the effect of erythromycin on gastric motor activity *in vivo* by evaluating gastric emptying. Intraperitoneal injection of erythromycin significantly promoted gastric emptying in hMTLR-Tg mice without any effect in WT mice. A previous study has shown that an erythromycin-derivative promoted gastric emptying in dogs [40]. Thus, our evidence suggests that hMTLR stimulation can promote gastric motor activity via a contractile response of smooth muscles in hMTLR-Tg mice. For future studies, we plan to investigate the prokinetic effect of human motilin and erythromycin in pathological conditions. Going forward, the hMTLR-Tg model will be a useful tool to study the interactions between the motilin and ghrelin pathways.

## Conclusions

In the stomach of hMTLR-Tg mice, hMTLR was primarily expressed in smooth muscle and the myenteric plexus, but in the latter structure, not colocalized with cholinergic neurons. Human motilin and erythromycin promoted a gastric motor response both *in vivo* and *in vitro* conditions. This action is mediated by the direct contraction of smooth muscle via influx of extracellular Ca^2+^. Thus, hMTLR-Tg mice may be a useful model for the evaluation of MTLR agonists as gastric prokinetic agents.

## Author Contributions

Conceived and designed the experiments: SK BM. Performed the experiments: SK AT MS AY KM. Analyzed the data: SK KM BM. Contributed reagents/materials/analysis tools: SK KM BM. Wrote the paper: SK. Checked and revised the manuscript: BM.

